# Expecting the unexpected: Great apes anticipate surprise in others

**DOI:** 10.64898/2026.10.12.758475

**Authors:** I.B. Laumer, L.S. Lewis, J. Eckert, S. Winkler, M. Köster, R. Mundry, D. Haun, C. Krupenye, E.A. Cartmill

**Author notes:** Shared 1^st^ authorship (I.B. Laumer, L.S. Lewis). Shared last authorship (C. Krupenye, E.A. Cartmill).

## Abstract

Humans adeptly navigate complex social environments by attempting to anticipate the actions and emotional responses of others. Although our closest ape relatives have sophisticated social cognitive abilities, we do not know whether they have a cognitive understanding of others’ emotions. For the epistemic emotion surprise, a cognitive understanding would require sensitivity to an agent’s belief, to reality, and to any misalignment between the two – allowing one to model the mental conditions that drive surprise. In a non-invasive eye tracking study with 31 chimpanzees and bonobos, apes showed evidence of a cognitive understanding of surprise: they detected the misalignment between an agent’s belief and reality, exhibiting greater pupil dilation right before an agent encountered an unexpected outcome relative to an expected one. This suggests that apes and humans share a more complex understanding of others’ mental lives than previously thought, capacities that likely evolved millions of years ago in a common ancestor.

## Introduction

Social animals, including humans and most non-human primates, navigate complex social environments in which anticipating the behavior and emotional reactions of others offers critical adaptive benefits (Frith & Frith, 2003; Genschow et al., 2018). Predicting how others will react or feel during social interactions could help individuals to build and manage relationships, avoid conflicts, and identify opportunities for learning. Yet, despite decades of research, we know surprisingly little about animals’ understanding of others’ emotions (Ferretti & Papaleo, 2019). Our closest ape relatives, for example, can distinguish and even categorize affective signals (e.g., matching scream faces to scream vocalizations in a touchscreen task: (Parr, 2001, 2004; Parr et al., 1998) and respond to such signals in natural contexts (e.g., consoling others in distress: (Clay & De Waal, 2013; De Waal & van Roosmalen, 1979). But what conception do they hold of individual affective states beyond an association between co-occurring outward signals, like facial and vocal expressions? Do they have a cognitive understanding, or mental model, of any emotion?

Although this question has often seemed empirically intractable, surprise offers a powerful inroad for understanding whether animals represent any emotions as internal states. Surprise is an epistemic emotion that arises when an agent’s expectations are violated (Reisenzein et al., 2019), sometimes referred to expectation violation (Afifi & Metts, 1998). This occurs when the agent has an inner model of the world that does not conform to reality. When the agent confronts the true state of the world, and the misalignment between their beliefs and reality, they experience surprise. For example, an agent will be surprised when someone jumps out of the closet because she believes that the closet is empty and therefore does not expect anyone to jump out of it. For a bystander to hold a mental model of another agent’s surprise, they must be sensitive to the agent’s epistemic state (i.e., belief), to reality, and to any misalignment between the two. Sensitivity to this misalignment offers an untapped marker that a bystander (human and non-human animals) may hold a mental model of surprise. And importantly, because the bystander’s mental model allows surprise to be *anticipated*, this marker can be measured before the agent produces any outward signals of surprise (e.g., facial or vocal expressions), precluding the bystander from simply responding to behavioral cues associated with the internal state of surprise.

Because surprise results from the violation of an agent’s epistemic states (i.e., beliefs), anticipating surprise recruits theory of mind abilities to infer others’ mental states (Krupenye & Call, 2019; Premack & Woodruff, 1978). Indeed, in a now classic commentary that inspired the false belief test, Dennett (Dennett, 1978) argued that this marker of a cognitive understanding of surprise, detecting misalignment between another agent’s beliefs and reality, would also be diagnostic evidence for false belief reasoning: “Very young children watching a Punch and Judy show squeal in anticipatory delight as Punch prepares to throw the box over the cliff. Why? Because *they know Punch thinks Judy is still in the box*. They know better; they saw Judy escape while Punch’s back was turned. We take the children’s excitement as overwhelmingly good evidence that they understand the situation - they understand that Punch is acting on a mistaken belief.” In a pioneering set of experimental studies that built on this anecdote, Moll and colleagues (Moll et al., 2016; Moll et al., 2017) demonstrated that young human children under the age of three indeed show this anticipatory marker of surprise understanding. As an agent approached a box, children showed more expressions of suspense or tension (e.g., biting their lip) when the agent’s belief about the contents of the box was misaligned with reality (i.e., when it was false) than when their belief aligned with reality.

There is evidence that a cognitive understanding of others’ surprise thus emerges early in human ontogeny. This ability to recognize and project others’ expectations about the world is central to a diversity of deeply human behaviors, from complex forms of coordination to crafting a joke that others will find funny. But what are the evolutionary origins of this fundamental human capacity? As Dennett (Dennett, 1978) asked nearly fifty years ago, “Would chimpanzees exhibit similar excitement if presented with a similar bit of play acting (in a drama that spoke directly to their “interests”)?”

The present study aims to test this question. Chimpanzees and bonobos are humans’ closest relatives, and they exhibit a suite of complex social behaviors and cognitive abilities that suggest they may share with humans a cognitive understanding of surprise. First, all great ape species engage in playful teasing, which involves intentional, one-sided, repeated and often boundary-pushing behaviors designed to provoke a reaction in the target ((Eckert et al., 2020; Laumer et al., 2024), see also (Krupenye & Hare, 2018)). Playful teasing often involves the violation of others’ expectations, such as offer-and-withdrawal games or sudden jumps. These playful teasing actions suggest that great apes may be able to anticipate that the target has formed a specific expectation (e.g., to receive an object) and may act with the intention of violating that expectation in order to provoke surprise (Eckert et al., 2020).

Second, apes have shown nuanced attention to their social worlds (e.g., (Carvajal & Krupenye, 2025; Cheney & Seyfarth, 1980; Krupenye & Hare, 2018; Lewis et al., 2021; Lewis et al., 2025; Lewis et al., 2023; Wittig et al., 2014)), a capacity to predict others’ behavior across multiple contexts (e.g., (Kano & Call, 2014; Kano & Hirata, 2015; Myowa-Yamakoshi et al., 2012)) and a sensitivity to others’ epistemic states (e.g., (Crockford et al., 2012; Girard-Buttoz et al., 2020; Townrow & Krupenye, 2025)), including their beliefs (Buttelmann et al., 2017; Kano et al., 2019; Krupenye et al., 2016; Lurz et al., 2022; Padberg et al., 2025). Critically, however, to our knowledge no studies have investigated whether apes (or any other non-human animals) show sensitivity to the misalignment between an agent’s beliefs and reality ahead of a surprising event. That is, no studies have directly tested whether any animal has a cognitive understanding of surprise.

To address this important gap, we presented chimpanzees and bonobos with video scenarios in which a protagonist held either a true or false belief about the contents of a box. We used an eye-tracker (Hopper et al., 2021; Lewis & Krupenye, 2022) to non-invasively record apes’ gaze and changes in pupil size, as they watched these videos. Because the pupils of both non-human apes and humans dilate in response to sympathetic activity of the autonomic nervous system (Hepach & Westermann, 2016; Joshi & Gold, 2020; Kret et al., 2014), under controlled lighting conditions, pupil dilation provides a validated measure of arousal (e.g., (Bradley et al., 2008; Hepach et al., 2021)). We predicted that apes would exhibit greater arousal (i.e., increased pupil dilation) as the agent approached the box with a false belief about its contents than with a true belief about its contents. Such a finding would indicate that apes could detect the misalignment between the agent’s belief and reality ahead of a surprising event, and therefore that they possess some cognitive understanding of surprise. We also investigated attention to the agent’s face and body during their approach as a further measure of surprise anticipation, although apes are known to exhibit much less attention to the faces of goal-directed agents than human participants (e.g., (Myowa-Yamakoshi et al., 2012)). Finding evidence for anticipation of others’ surprise in both chimpanzees and bonobos would suggest that this capacity was likely present in our last common ancestor, shedding light on the evolutionary origins of emotional and cognitive perspective-taking.

## Methods

### Participant information

A total of 31 apes participated in this study: nine chimpanzees (*Pan troglodytes*, three females and six males) living at the Edinburgh Zoo, and 14 chimpanzees (nine females and five males) and eight bonobos (*Pan paniscus*, four females and four males) living at the Leipzig Zoo (see SI, section A, Table S1). Apes were housed in social groups and participated in all experiments on a voluntary basis (for housing information see SI, section A).

### Ethics

Experimental protocols were approved by the School of Psychology and Neuroscience Animal Ethics Committee at the University of St. Andrews and the ethics committee of the Max Planck Institute for Evolutionary Anthropology, Zoo Leipzig. Participants were tested in the testing rooms prepared for each species. Apes’ daily participation in this study was completely voluntary. They received regular feedings and daily enrichment and had *ad libitum* access to water. Animal husbandry and research protocol complied with international standards (the Weatherall report, The use of non-human primates in research) and institutional guidelines (EAZA minimum standards for the accommodation and care of animals in zoos and aquaria; WAZA Ethical guidelines for the conduct of research on animals by zoos and aquariums; Guidelines for the treatment of animals in behavioral research and teaching (ASAB/ABS)). The research adhered to all German and Scottish laws regarding animal holding.

### Research design and video stimuli

The video stimuli consisted of four unique stories, each with a true belief and a false belief version (i.e., eight test sequences total). Each story involved a protagonist hiding an object in a box and leaving, an antagonist appearing and changing the location, identity, or integrity of the object while the protagonist was either fully absent (false belief condition) or covertly watching (true belief condition). Finally, at the end of the video, the protagonist returns and approaches the box to retrieve the object. Thus, exclusively in the false belief condition, the protagonist’s belief was misaligned with reality and they were poised to be surprised by the contents of the box, which had been altered in their absence. Two stories were designed to have exciting outcomes (the addition of a snake and a hidden opponent with a weapon), while the other two stories were designed to have less exciting outcomes (a toy disappeared), and a tool was destroyed). Each video was immediately preceded by a familiarization video that depicted the protagonist’s interest in the target object, the antagonistic relationship between the protagonist and antagonist, and the setting (including the protagonist hiding and retrieving the object from the box). The entire duration of the familiarization phase and the test phase (true or false belief version) was kept under one minute to maximize apes’ attention across the entire stimuli. Videos and audio were edited using DaVinci Resolve (version 17) and iMovie (version 10.2.4). We used the same lighting across conditions within each story to avoid a potential impact on pupillometry measures. To minimize variation in local luminance, luminance of the left side was matched to the luminance on the right side of each frame, by applying a linear gradient using Matlab (version 2021a). The video clips that were edited and combined to make the true and false belief versions of each scene were identical except for the brief section in the middle in which the protagonist either watched or did not watch the transformation.

### Procedure

Apes were shown video stimuli using Tobii eye-tracking hardware and software (Tobii X120, Tobii Pro Studio, Tobii AB, Sweden). To encourage the apes to sit still in front of the screen and eye tracker, they could drink diluted juice from a straw while their eye movements and pupil dilation were non-invasively measured (see Figure 1). Before testing, we conducted a two-point automated calibration for each ape subject by presenting a small video clip on each reference point (see SI, section B for details). Each subject’s unique calibration was used throughout the entire testing period. Prior to the start of every session, the calibration accuracy was manually evaluated by presenting subjects with a grid image containing several objects and observing whether gaze was mapped accurately onto those objects (Figure S1). Calibrations were repeated when necessary. Calibration errors are typically less than a degree with this procedure (Hopper et al., 2021).

**Figure 1.**
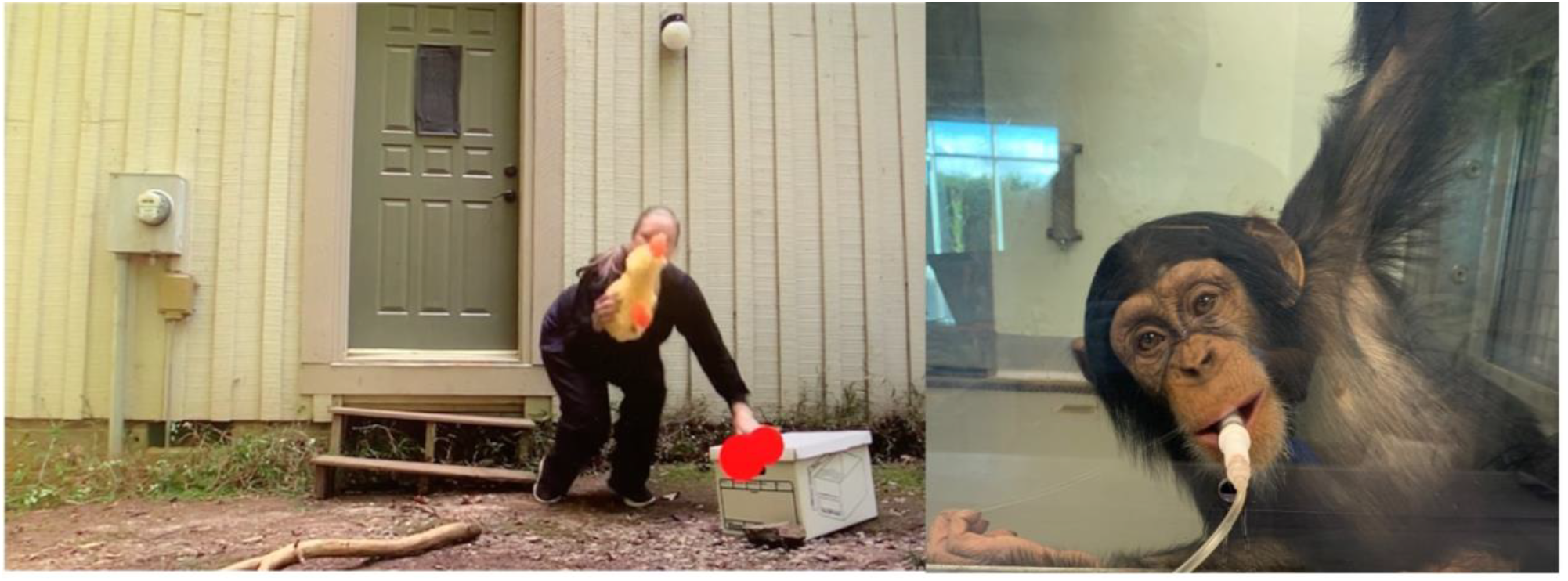
Eye-tracking setup and frame depicting subject’s gaze. **Left:** The screenshot shows the moment before the protagonist hides the toy in the box (snake story; author EAC is depicted here). The red dots depict the ape’s eye gaze, indicating where they are looking on the screen (in this case at the box). **Right:** A chimpanzee (from Edinburgh Zoo) drinking diluted fruit juice from a straw while watching a test video on a monitor through a polycarbonate panel (plastic material used for eye tracking with apes). Apes voluntarily participated and could choose to walk away from the eye tracking set up at any time.

Once the ape was attending to the screen, the experimenter started the familiarization video. The test video was presented directly after the familiarization video, once the ape was again attending to the screen. Apes watched one familiarization and test sequence per day and saw all eight test videos across eight separate test days in a semi-randomized order. Stimuli alternated between true and false belief conditions across test days, and apes never saw the two (true and false belief) videos from the same story on consecutive test days. Half of the participants started with a true belief condition, and half started with a false belief condition.

### Metrics and Statistical Analyses

To calculate our primary dependent measure, pupil dilation, for each trial we computed a subject’s average pupil diameter during fixations to the entire screen, separately for the baseline phase (a 2s time window, that was considered to be not exciting, in which there were minimal movements and actions on the screen) and the anticipatory phase (beginning when the protagonist began their final approach to the box and ending when the protagonist began to lift the lid of the box; the video stopped before the box’s contents were visible; for an overview of the phases see Table S2). Pupil measurements were retained only for frames for which the eyes were flagged as valid by the eye tracker. For the Leipzig data, the Tobii software automatically provided a validity label to the data export, and only samples labelled “valid” were retained. For the Edinburgh data, validity was automatically coded numerically (0-4), and codes 0 and 1 (highly confident to confident) were treated as valid. Pupil measurements for any eyes that did not meet these criteria were excluded. Pupil dilation was then calculated by subtracting average baseline pupil diameter from average anticipatory phase pupil diameter. To explore gaze duration patterns, we also calculated total fixation duration during the anticipatory phase for two static rectangular areas-of-interest (AOIs): one encompassing just the box (box AOI) and a second encompassing the protagonist, her path toward the box, and the box itself (protagonist AOI). We quantified gaze allocation as the proportion of total fixation time directed toward each AOI during the anticipatory window, calculated by dividing fixation duration within the AOI by total fixation duration to the entire screen during the anticipatory window. Fixations were identified using Tobii’s default fixation filter (Tobii I-VT filter). In the gaze duration models, observations were weighted by total looking time, as proportions are more precise when based on larger amounts of looking time. This normalized measure captures the relative allocation of gaze to the AOI while accounting for differences in the total looking time across trials.

To investigate whether condition (true or false belief) predicted pupil dilation or gaze duration, we fitted Linear Mixed Models (LMM) and two Generalized Linear Mixed Models (GLMM; (Baayen, 2008)) in R (version 4.4.2; (R Core Team, 2024)) (Table S3). The models’ fixed effects were condition, species, story, all their interactions up to third order, trial order, and location (Edinburgh or Leipzig). Originally, we also planned to control for luminosity; however, including the interaction between condition, species, and story, up to third order, and luminosity was not possible because for any given combination of condition and story luminosity was a constant. Therefore, we fitted a model including condition, species and story and included the two two-way interactions between species on the one hand and condition and story on the other, to check for evidence that luminosity affected pupil dilation in this model. However, this did not seem to be the case (estimate±SE=0.13±0.83, t=0.16, df=57.3, p=0.88). Therefore, we did not control for luminosity in our main model of pupil dilation. All models included subject ID as a random intercept. To avoid an overconfident model and keep type I error rate at the nominal level of 0.05, we also included all theoretically identifiable random slopes (Barr et al., 2013; Schielzeth & Forstmeier, 2009), namely those of condition, story, and trial order. Originally, we included parameters for the correlations among the random intercept and slopes. However, due to convergence and identifiability problems, we had to exclude these correlation parameters from all models. In the gaze duration models, random slope variance was negligible or convergence could not be achieved even without estimated correlations; we therefore further simplified these models to a random intercept of participant only. As an overall test of the effect of condition, our key predictor of interest, we compared each full model to a respective null model that was identical to the full model except for the removal of condition and its interactions from the fixed effects part. Such a full-null model comparison aims at avoiding cryptic multiple testing (Forstmeier & Schielzeth, 2011).

We fitted the model with pupil dilation difference as the dependent variable with an LMM with Gaussian error distribution and identity link function (McCullagh & Nelder, 1989). This model was fitted using the function lmer of the package lme4 (version 1.1-35.5; (Bates, 2015). The response variable in the gaze duration models, the raw proportion of looking time during the test phase, was bound between 0 and 1 and fitted using the function glmmTMB of the equally named package (version 1.1.10; (Brooks et al., 2017)) using a beta error distribution (Bolker, 2008) and logit link function. Prior to fitting the models, we transformed the response to avoid values being exactly zero or one (Smithson & Verkuilen, 2006). For all models, we also z-transformed trial order to a mean of zero and a standard deviation of one to achieve an easier interpretation of model estimates (Schielzeth, 2010) and ease model convergence. To include condition and story into the random effects part, we manually dummy coded and then centered them to a mean of zero.

The p-values we report for the gaze duration models are based on likelihood ratio tests (Dobson, 2002). For these models we estimated the p-value of individual terms by dropping them from the model, one at a time, and comparing the simpler models with the respective full model (R function drop1). The full-null model comparisons for all models also utilized a likelihood ratio tests. However, for the gaze duration models, we determined the significance of individual predictors by means of the Satterthwaite approximation (Luke, 2017) as implemented in the function lmer of the package lmerTest (version 3.1-3; (Kuznetsova et al., 2017). If an interaction was not significant in a given model, we removed it from the model to obtain unconditional estimates (Aiken, 1991) of the respective lower order terms. We determined model stability by dropping individual apes from the data, one at a time, and comparing the estimates derived for the models fitted with the respective subsets with those obtained for the respective full data set (Nieuwenhuis et al., 2012). This revealed all models to be of moderate to excellent stability (see minimum and maximum estimates in the tables S4, S6, S7). We obtained confidence intervals of model estimates and potentially fitted values by means of parametric bootstraps (N=1,000 bootstraps; functions bootMer of the package lme4 or simulate of the package glmmTMB).

In the gaze duration models, the response was not overdispersed for the protagoinst AOI (dispersion parameter: 1.01) but showed indication of overdispersion for the box AOI (dispersion parameter: -1.43); results from this model should therefore be interpreted with caution. The sample analyzed with the models encompassed 206 gaze durations (one for each trial in which gaze duration in the baseline and test phase could be measured) of 31 apes (8 bonobos and 23 chimpanzees) or 140 pupil dilation differences of 30 apes (8 bonobos and 22 chimpanzees; for an overview see SI, Table S10; note that we took a conservative approach by excluding all measurements for which the eye tracker was uncertain about whether the data truly originated from the left (or right) eye, or for which the eye was not detected).

## Results

As predicted, condition had a clear impact on pupil diameter (full-null model comparison: χ2 = 19.402, df = 8, P = 0.013), but the interaction between condition, species, and story was not significant (F(3, 50.2) = 1.401, P = 0.254; see SI, section C1, Table S4). After the removal of this non-significant interaction, we found a significant effect of the interaction between condition and story (F(3, 55.7) = 3.651, P = 0.018; SI, section C1, Table S5). As expected, we found larger pupil dilations in the false than the true belief condition in the toy, snake, and to a lesser extent tool story, though not in the weapon story (Figure 2).

**Figure 2.**
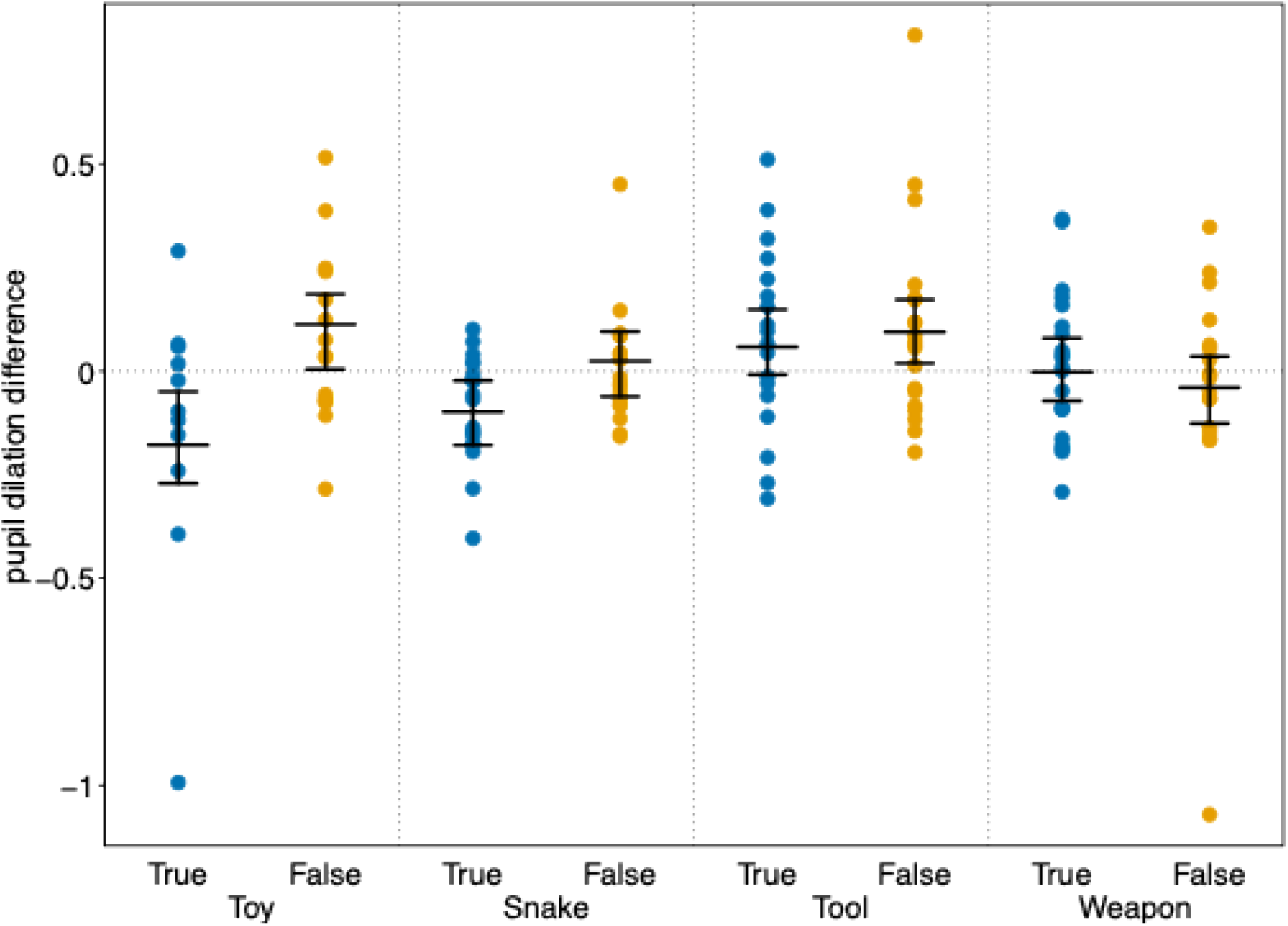
Difference in pupil dilation (test phase minus baseline phase) in the true and false belief condition of each video story. Horizontal lines with error bars indicate the fitted model and its 95% confidence limits with all other fixed effects terms in the model being at their average. Dots depict the individual pupil dilation differences.

Overall, in the false belief condition, pupil diameter was typically larger in the test window than in the baseline window (mean = 0.036 mm, SD = 0.230 mm, n = 71). In the true belief condition, pupil diameter was typically smaller in the test window than in the baseline window (mean = −0.025 mm, SD = 0.221 mm, n = 69).

We found no evidence for a condition effect on anticipatory attention (i.e., gaze duration). For the protagonist AOI, the full-null model comparison showed no evidence that condition, or its interactions with species and story, affected proportion of looking time (χ^2^ = 2.57, df=8, P = 0.958; see SI, section C2, Table S6). For the box AOI, the full-null model comparison likewise showed no evidence for an effect of condition or its interactions (χ^2^ = 2.44, df=8, P = 0.965; see SI, section C2, Table S7), although this model showed indication of overdispersion and the results should be interpreted with caution.

## Discussion

Our findings suggest that our closest living relatives, chimpanzees and bonobos, possess some cognitive understanding of surprise. Apes demonstrated higher arousal, as measured by larger pupil dilation, before a protagonist discovered an unexpected outcome compared to an expected one. This result implies that apes were sensitive to both the agent’s belief and reality and were able to detect the misalignment between the two, leading them to experience anticipatory arousal in advance of the agent’s surprise.

Critically, because apes’ arousal was anticipatory, it occurred before the agent discovered the unexpected state of reality and therefore before the agent produced any outward cues to her impending surprise. Thus, there were no associative cues, like facial or vocal expressions of surprise, available for apes to respond to. Indeed, the videos ended just as the protagonist started to lift the lid of the box and before they were able to see the change. Importantly, the true and false belief versions of each video story, including the anticipatory phase, used identical video footage, except for the brief central manipulation that determined the protagonist’s belief (covertly witnessing the antagonist’s key action with the box in the true belief condition but not in the false belief condition).

We did not find a significant effect of looking time towards the protagonist or the box, which we included as an additional measure. One possible explanation is that great apes are known to exhibit less attention to the faces of agents than do human participants (Myowa-Yamakoshi et al., 2012). As we found a significant difference in pupil size between conditions, another possibility is that the apes may have grasped the relevant information quickly (apes show high cognitive processing speed (Inoue & Matsuzawa, 2007), but see: (Silberberg & Kearns, 2009)) and generally looked at the protagonist only briefly, so that the measure was thus less sensitive to differences between conditions. In addition, the model analyzing looking time towards the box showed signs of overdispersion, so this result should be interpreted with caution.

For over 50 years, researchers have wondered about the mentalizing capacities of our closest living phylogenetic cousins. In a 1978 commentary on Premack and Woodruff’s seminal *Behavioral and Brain Sciences* paper, “Does the chimpanzee have a theory of mind?” (Premack & Woodruff, 1978); Dennett provided theoretical and practical foundations for exploring whether great apes have a theory of mind by testing whether they experience anticipatory arousal just before a mistaken agent is about to be surprised. By showing apes dynamically social movie stories (as suggested by: (Kano et al., 2020), we were able to create, as Dennett (Dennett, 1978) put it, “drama[s] that spoke directly to their ‘interests’. In three of the four stories, we found Dennett’s predicted effect: greater arousal in the false belief condition. It is unclear why the weapon story did not elicit any pupillary difference. It is possible that, because the weapon story included the aggressive antagonist hiding in the box in the anticipatory phase, the apes’ excitement remained high even in the true belief condition, where they may have anticipated a conflict between the agents. Since it was the only condition where the antagonist remained in the scene, albeit hidden from view, the apes may have mentalized the antagonist, reducing their focus on the protagonist’s expectations. Nonetheless, the findings of the other stories suggest that apes can anticipate surprise across a diversity of contexts.

We obtained the most pronounced effects in one of the less exciting stories, the toy story, and one of the more exciting stories, the snake story. These scenarios involved straightforward change-of-content manipulations to the box. However, they differed in whether an object was removed (toy story) or added (snake story), suggesting that apes can represent both kinds of false beliefs. Our study thus adds to a growing body of work demonstrating apes’ sensitivity to others’ mental states, including beliefs, across varied contexts and via an increasing diversity of measures. Past research demonstrated that apes can predict the behavior of an agent with a false belief (Kano et al., 2019; Krupenye et al., 2016), but this is the first study to show that apes exhibit anticipatory arousal when an agent’s belief does not conform to reality—just like young human children (Moll et al., 2016; Moll et al., 2017; Scott, 2017). Anticipatory arousal, especially in benign scenarios like the toy story, suggests that apes not only computed the agent’s belief but also detected its misalignment with reality. This study therefore highlights that tracking others’ epistemic states, apes may be evaluating the discrepancy between an agent’s belief and the ape’s own knowledge of the situation, in contrast to prominent accounts that only humans compare their mental state ascriptions with their representation of reality (Tomasello, 2018).

Our finding that apes appear to detect the misalignment between another agent’s expectations and reality suggest that they possess a cognitive understanding of surprise and a mental model of at least one internal emotional state. This form of perspective-taking, a key component of empathy (De Waal & Preston, 2017), is therefore within the cognitive potential of our closest living relatives and likely evolved at least 6-9 million years ago in a common evolutionary ancestor. That apes seem to understand the mental causes of surprise also raise the possibility that they may be capable of grasping the causes of other more basic emotions, spotlighting an important target for future research. Future work should also investigate other aspects of apes’ mental model of surprise, such as whether they can use an agent’s surprise response to infer the identity of the object that caused it (Wu et al., 2024; Wu et al., 2021).

Our findings also shed light on apes’ fascinating propensity for playfully teasing one another (Laumer et al., 2024). Playful teasing, found in all four great ape species, often involves provoking surprise and our findings support the theory that, in these situations, apes may be monitoring their partner’s expectations and intentionally violating them (Eckert et al., 2020; Winkler & Cartmill, 2026). Investigating surprise anticipation in the other two great ape species (gorillas and orangutans) would help clarify whether these mechanisms are a cognitive prerequisite of teasing and present in all species that playfully tease. Our discovery that apes possess some cognitive understanding of surprise suggests that they share with humans a more complex understanding of others’ mental lives than previously thought, and that the underlying cognitive mechanisms are evolutionarily ancient features of the human mind.

## Author contributions

IBL, LSL, CK, JE, SW, and EAC designed the study. IBL, EAC, SW, CK, and JE filmed the video stimuli, and IBL, EAC, and MK edited and created the video stimuli. LSL collected the data. MK provided input and IBL and RM analyzed the data. IBL, LSL, and CK wrote the manuscript, and JE, SW, DH, MK, RM, and EAC edited the paper.

## Funding

This project was funded by a Templeton World Charity Foundation grant (grant no. TWCF0318) to E.A.C. S.L.W. is supported by the NSF Graduate Research Fellowship Program (grant no. DGE-2034835).

### Acknowledgements

We are grateful to all of the chimpanzees and bonobos who participated in this study. This project was funded by a Templeton World Charity Foundation grant (TWCF0318) to Erica A. Cartmill. We thank Philipp Gresser and Dr. Christina Gresser, and Kaye Brown Cartmill for their acting performance in filming the video stimuli and Wolfgang Laumer and Thomas Bildl for dubbing the sound. We would like to thank Edinburgh Zoo and Leipzig Zoo for permission and support in conducting this research. At Edinburgh Zoo, we thank Kate Grounds, Donald Gow, Roslin Talbot, Callum Gibson, and the Budongo Research Unit keepers. At the Max Planck Institute for Evolutionary Anthropology and Leipzig Zoo, we thank Hanna Petschauer, Daniel Hanus, Daniel Geissler, and the other animal caregivers and staff. We are grateful to the Royal Zoological Society of Scotland (RZSS) for core financial support to the RZSS Edinburgh Zoo’s Budongo Research Unit and the Max Planck Society for core financial support to the Wolfgang Köhler Primate Research Center at Zoo Leipzig, where this project was carried out.

## Data Availability

Data and analysis scripts will be uploaded during the submission process and upon publication the data will be shared publicly as supplementary materials and on Dryad.

