## Supplementary Material for "Expecting the unexpected: Great apes anticipate surprise in others"

##### A) Subject information and housing conditions

**Table S1** Species, names, sex, age, and zoos the apes were housed in.

| Participant | Species | Sex | Date of Birth | Age at Testing | Facility |
| --- | --- | --- | --- | --- | --- |
| Frek | Chimpanzee | M | 10/21/93 | 28.05 | Edinburgh Zoo |
| Liberius | Chimpanzee | M | 1/20/99 | 22.79 | Edinburgh Zoo |
| Louis | Chimpanzee | M | 7/26/76 | 45.29 | Edinburgh Zoo |
| Qafzeh | Chimpanzee | M | 3/31/92 | 29.61 | Edinburgh Zoo |
| Rene | Chimpanzee | M | 2/21/93 | 28.71 | Edinburgh Zoo |
| Velu | Chimpanzee | M | 6/24/14 | 7.36 | Edinburgh Zoo |
| Edith | Chimpanzee | F | 4/11/96 | 25.58 | Edinburgh Zoo |
| Eva | Chimpanzee | F | 12/9/80 | 40.92 | Edinburgh Zoo |
| Kilimi | Chimpanzee | F | 2/20/93 | 28.72 | Edinburgh Zoo |
| Amira | Bonobo | F | 5/15/17 | 5.30 | Leipzig Zoo |
| Jasongo | Bonobo | M | 8/2/90 | 32.10 | Leipzig Zoo |
| Kuno | Bonobo | M | 11/26/96 | 25.78 | Leipzig Zoo |
| Lexi | Bonobo | F | 9/13/99 | 22.98 | Leipzig Zoo |
| Luiza | Bonobo | F | 1/27/05 | 17.60 | Leipzig Zoo |
| Tayo | Bonobo | M | 7/22/16 | 6.11 | Leipzig Zoo |
| Yaro | Bonobo | M | 4/18/13 | 9.37 | Leipzig Zoo |
| Yasa | Bonobo | F | 8/27/97 | 25.02 | Leipzig Zoo |
| Azibo | Chimpanzee | M | 4/14/15 | 7.38 | Leipzig Zoo |
| Changa | Chimpanzee | F | 3/3/11 | 11.50 | Leipzig Zoo |
| Corrie | Chimpanzee | F | 12/12/76 | 45.75 | Leipzig Zoo |
| Frank | Chimpanzee | M | 12/25/19 | 2.68 | Leipzig Zoo |
| Frodo | Chimpanzee | M | 11/28/93 | 28.77 | Leipzig Zoo |
| Kisha | Chimpanzee | F | 3/4/04 | 18.50 | Leipzig Zoo |
| Makeni | Chimpanzee | M | 3/14/18 | 4.47 | Leipzig Zoo |
| Natascha | Chimpanzee | F | 3/28/80 | 42.45 | Leipzig Zoo |
| Ohini | Chimpanzee | M | 3/25/16 | 6.44 | Leipzig Zoo |
| Riet | Chimpanzee | F | 11/11/77 | 44.83 | Leipzig Zoo |
| Sandra | Chimpanzee | F | 6/9/93 | 29.24 | Leipzig Zoo |
| Swela | Chimpanzee | F | 10/19/95 | 26.88 | Leipzig Zoo |
| Tai | Chimpanzee | F | 8/12/02 | 20.06 | Leipzig Zoo |
| Youma | Chimpanzee | F | 3/25/18 | 4.44 | Leipzig Zoo |

#### Housing conditions

Each of the ape groups at Leipzig Zoo in Germany and at the Edinburgh Zoo in Scotland were housed in large indoor and outdoor enclosures, which were equipped with various climbing structures and enrichment devices such as shaking and raking boxes. All apes in this study were fed several times a day with a diet consisting of a varying mixture of vegetables, greens, fruits, primate pellets, seeds. Water was available ad libitum. Additionally, apes received daily enrichment such as burlap sacks and cardboard, often filled with food.

#### B) Additional information

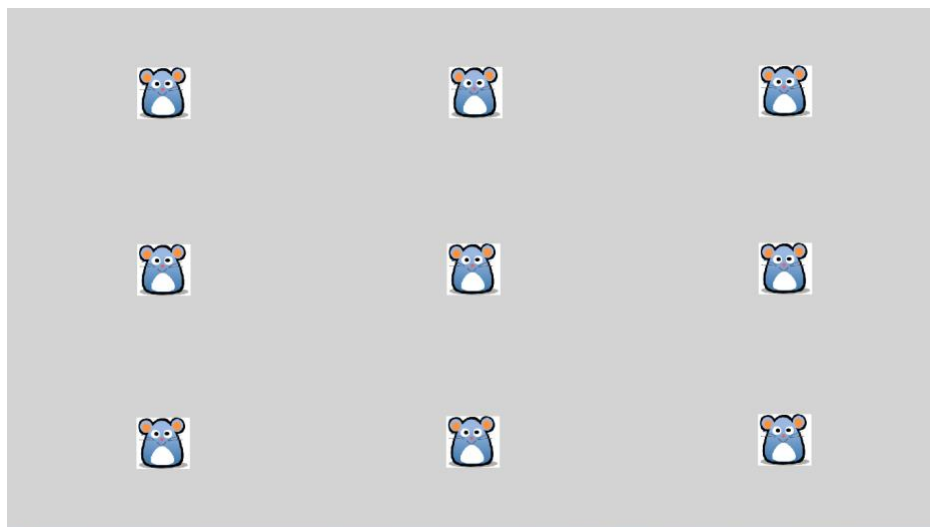

**Figure S2** Screenshot of the mouse grid to check calibration accuracy. This calibration screen was presented directly before each familiarization trial to manually check whether calibrations were accurate before playing the familiarization and test videos. Familiarization and test videos were only shown after determining that the calibration was accurate.

**Table S2** Duration of video stimuli and timing of baseline and test measurements.

|  | Duration familiarization video | Duration true belief story | Duration false belief story | Baseline measurement | Test measurement |
| --- | --- | --- | --- | --- | --- |
| **Toy** | 0:27:22 | 0:33:03 | 0:33:03 | 0:10:23 to 0:12:23 | 0:26:20 to 0:33:03 |
| **Tool** | 0:29:23 | 0:44:05 | 0:44:05 | 0:16:08 to 0:18:08 | 0:38:11 to 0:44:05 |
| **Snake** | 0:34:08 | 0:34:18 | 0:34:18 | 0:07:00 to 0:09:00 | 0:29:11 to 0:34:18 |
| **Weapon** | 0:22:17 | 0:40:18 | 0:40:18 | 0:04:06 to 0:06:06 | 0:33:09 to 0:40:18 |

**Table S3** Model structure for each model.

| model | full model structure |
| --- | --- |
| 1) Pupil dilation difference | pupil dilation difference ~ condition*species*story + order + location + (1 + condition true + tool story + toy story + weapon story + order \| \|name of participant), REML=F |
| 2) Proportion looking time difference at box area | Proportion looking time at box ~ condition*species*story + order + location + (1 \|name of participant), family=beta |
| 3) Proportion looking time at protagonist area including box | Proportion looking time at protagonist ~ condition*species*story + order + location + (1 + condition true + tool story + toy story + weapon story + order \| \|name of participant), family=beta |

Note that order was z-transformed to a mean of zero and a standard deviation of 1 and that condition true, tool story, toy story and weapon story are centered dummy variables.

### C) Additional statistical results

#### C1) Pupil dilation

**Table S4** Results of the full model (including the three-way interaction between condition, species, and story) with pupil dilation as the dependent measure.

| term | Estimate | SE | CIlwr | CIupr | F | NumDF | DenDF | P | min | max |
| --- | --- | --- | --- | --- | --- | --- | --- | --- | --- | --- |
| (Intercept) | 0.032 | 0.081 | -0.110 | 0.166 |  |  |  |  | -0.027 | 0.065 |
| conditionTrue | -0.146 | 0.090 | -0.318 | 0.027 |  |  |  |  | -0.176 | -0.096 |
| speciesChimp | -0.065 | 0.090 | -0.226 | 0.099 |  |  |  |  | -0.098 | -0.001 |
| storyTool | -0.032 | 0.099 | -0.212 | 0.158 |  |  |  |  | -0.084 | 0.033 |
| storyToy | 0.044 | 0.129 | -0.143 | 0.233 |  |  |  |  | -0.074 | 0.144 |
| storyWeapon | -0.246 | 0.105 | -0.431 | -0.078 |  |  |  |  | -0.301 | -0.083 |
| z.order | 0.017 | 0.016 | -0.010 | 0.046 | 0.997 | 1 | 22.837 | 0.328 | 0.001 | 0.030 |
| locationLeipzig | 0.036 | 0.043 | -0.040 | 0.110 | 0.620 | 1 | 22.982 | 0.439 | 0.006 | 0.067 |
| conditionTrue:speciesChimp | 0.042 | 0.116 | -0.184 | 0.266 |  |  |  |  | -0.011 | 0.077 |
| conditionTrue:storyTool | 0.191 | 0.128 | -0.092 | 0.444 |  |  |  |  | 0.074 | 0.236 |
| conditionTrue:storyToy | -0.289 | 0.143 | -0.548 | -0.006 |  |  |  |  | -0.438 | -0.162 |
| conditionTrue:storyWeapon | 0.246 | 0.126 | 0.023 | 0.527 |  |  |  |  | 0.111 | 0.341 |
| speciesChimp:storyTool | 0.164 | 0.122 | -0.079 | 0.382 |  |  |  |  | 0.095 | 0.215 |
| speciesChimp:storyToy | 0.062 | 0.160 | -0.195 | 0.288 |  |  |  |  | -0.055 | 0.177 |
| speciesChimp:storyWeapon | 0.298 | 0.134 | 0.070 | 0.528 |  |  |  |  | 0.140 | 0.353 |
| conditionTrue:speciesChimp:storyTool | -0.157 | 0.162 | -0.469 | 0.193 | 1.401 | 3 | 50.214 | 0.254 | -0.198 | -0.044 |
| conditionTrue:speciesChimp:storyToy | 0.231 | 0.191 | -0.109 | 0.603 |  |  |  |  | 0.104 | 0.370 |
| conditionTrue:speciesChimp:storyWeapon | -0.139 | 0.160 | -0.469 | 0.140 |  |  |  |  | -0.224 | -0.022 |

Indicated are fixed effect estimates together with their standard errors, 95% confidence limits, significance tests and the range of estimates (min and max) obtained when excluding levels of random effect factors one at a time. All factors were dummy coded; reference categories were False for condition, bonobo for species, snake for story, and Edinburgh for location.

Order was z-transformed to a mean of zero and a standard deviation (sd) of one, mean and sd of the original variable were 4.46 and 2.34.

**Table S5** Results of the reduced model (without the three-way interaction between condition, species, and story; three-way interaction removed, all two-way interactions kept) with pupil dilation as the dependent measure.

| term | Estimate | SE | CIlwr | CIupr | F | NumDF | DenDF | P |
| --- | --- | --- | --- | --- | --- | --- | --- | --- |
| (Intercept) | 0.013 | 0.077 | -0.117 | 0.150 |  |  |  |  |
| conditionTrue | -0.122 | 0.071 | -0.246 | -0.002 |  |  |  |  |
| speciesChimp | -0.040 | 0.076 | -0.171 | 0.079 |  |  |  |  |
| storyTool | 0.020 | 0.081 | -0.116 | 0.171 |  |  |  |  |
| storyToy | -0.006 | 0.123 | -0.173 | 0.159 |  |  |  |  |
| storyWeapon | -0.202 | 0.094 | -0.334 | -0.046 |  |  |  |  |
| z.order | 0.015 | 0.016 | -0.013 | 0.044 | 0.753 | 1 | 24.998 | 0.394 |
| locationLeipzig | 0.043 | 0.045 | -0.037 | 0.109 | 0.791 | 1 | 24.478 | 0.382 |
| conditionTrue:speciesChimp | -0.002 | 0.067 | -0.114 | 0.108 | 0.000 | 1 | 17.754 | 0.998 |
| conditionTrue:storyTool | 0.090 | 0.081 | -0.066 | 0.245 | 3.651 | 3 | 55.699 | 0.018 |
| conditionTrue:storyToy | -0.165 | 0.097 | -0.313 | 0.043 |  |  |  |  |
| conditionTrue:storyWeapon | 0.157 | 0.082 | 0.023 | 0.325 |  |  |  |  |
| speciesChimp:storyTool | 0.085 | 0.087 | -0.068 | 0.235 | 1.486 | 3 | 30.965 | 0.238 |
| speciesChimp:storyToy | 0.151 | 0.138 | -0.050 | 0.317 |  |  |  |  |
| speciesChimp:storyWeapon | 0.225 | 0.105 | 0.056 | 0.368 |  |  |  |  |

Indicated are fixed effect estimates together with their standard errors, 95% confidence limits and significance tests. All factors were dummy coded; reference categories were False for condition, bonobo for species, snake for story, and Edinburgh for location. Order was z-transformed to a mean of zero and a standard deviation (sd) of one, mean and sd of the original variable were 4.46 and 2.34.

### C2) Proportion looking time at protagonist AOI

**Table S6** Results of the full model with proportion looking time to the protagonist AOI as the dependent measure.

| term | Estimate | SE | CIlwr | CIupr | LRT | Df | P | min | max |
| --- | --- | --- | --- | --- | --- | --- | --- | --- | --- |
| (Intercept) | 0.556 | 0.525 | -0.462 | 1.562 |  |  |  | 0.324 | 0.885 |
| conditionTrue | 0.148 | 0.649 | -1.180 | 1.477 |  |  |  | -0.037 | 0.305 |
| speciesChimp | -0.310 | 0.574 | -1.467 | 0.772 |  |  |  | -0.645 | -0.069 |
| storyTool | -0.096 | 0.648 | -1.458 | 1.248 |  |  |  | -0.632 | 0.279 |
| storyToy | -0.633 | 0.655 | -1.875 | 0.677 |  |  |  | -0.845 | -0.339 |
| storyWeapon | -0.019 | 0.645 | -1.315 | 1.307 |  |  |  | -0.200 | 0.221 |
| z.order | 0.132 | 0.094 | -0.054 | 0.312 | 1.954 | 1 | 0.162 | 0.071 | 0.169 |
| locationLeipzig | -0.177 | 0.246 | -0.680 | 0.350 | 0.507 | 1 | 0.477 | -0.249 | -0.045 |
| conditionTrue:speciesChimp | -0.123 | 0.782 | -1.663 | 1.434 |  |  |  | -0.367 | 0.094 |
| conditionTrue:storyTool | 0.133 | 0.914 | -1.681 | 1.853 |  |  |  | -0.244 | 0.877 |
| conditionTrue:storyToy | -0.239 | 0.924 | -2.098 | 1.518 |  |  |  | -0.605 | -0.031 |
| conditionTrue:storyWeapon | -0.119 | 0.919 | -2.039 | 1.724 |  |  |  | -0.546 | 0.293 |
| speciesChimp:storyTool | -0.277 | 0.785 | -1.903 | 1.322 |  |  |  | -0.645 | 0.270 |
| speciesChimp:storyToy | 0.425 | 0.795 | -1.174 | 1.910 |  |  |  | 0.133 | 0.638 |
| speciesChimp:storyWeapon | -0.573 | 0.779 | -2.153 | 0.918 |  |  |  | -0.811 | -0.364 |
| conditionTrue:speciesChimp:storyTool | -0.304 | 1.101 | -2.550 | 1.918 | 0.893 | 3 | 0.827 | -1.105 | 0.058 |
| conditionTrue:speciesChimp:storyToy | 0.024 | 1.119 | -2.085 | 2.226 |  |  |  | -0.343 | 0.391 |
| conditionTrue:speciesChimp:storyWeapon | 0.695 | 1.098 | -1.562 | 2.918 |  |  |  | 0.297 | 1.130 |

Indicated are fixed effect estimates together with their standard errors, 95% confidence limits, significance tests and the range of estimates (min and max) obtained when excluding levels of random effect factors one at a time. All factors were dummy coded; reference categories were False for condition, bonobo for species, snake for story, and Edinburgh for location. Order was z-transformed to a mean of zero and a standard deviation (sd) of one, mean and sd of the original variable were 4.19 and 2.27.

#### C3) Proportion looking time at box AOI

**Table S7** Results of the full model with proportion looking time to the box as the dependent measure.

| term | Estimate | SE | CIlwr | CIupr | LRT | Df | P | min | max |
| --- | --- | --- | --- | --- | --- | --- | --- | --- | --- |
| (Intercept) | -2.819 | 0.406 | -3.667 | -1.978 |  |  |  | -2.966 | -2.764 |
| conditionTrue | 0.148 | 0.500 | -0.939 | 1.173 |  |  |  | 0.033 | 0.214 |
| speciesChimp | 0.152 | 0.436 | -0.787 | 0.990 |  |  |  | 0.069 | 0.272 |
| storyTool | 0.332 | 0.497 | -0.676 | 1.425 |  |  |  | 0.159 | 0.422 |
| storyToy | 0.267 | 0.499 | -0.811 | 1.243 |  |  |  | 0.083 | 0.469 |
| storyWeapon | 0.262 | 0.496 | -0.779 | 1.308 |  |  |  | 0.108 | 0.340 |
| z_order | 0.067 | 0.072 | -0.082 | 0.207 | 0.842 | 1 | 0.359 | 0.048 | 0.094 |
| locationLeipzig | -0.216 | 0.180 | -0.593 | 0.127 | 1.405 | 1 | 0.236 | -0.270 | -0.114 |
| conditionTrue:speciesChimp | -0.353 | 0.603 | -1.609 | 0.879 |  |  |  | -0.446 | -0.238 |
| conditionTrue:storyTool | -0.447 | 0.705 | -1.967 | 0.902 |  |  |  | -0.682 | -0.192 |
| conditionTrue:storyToy | -0.054 | 0.708 | -1.495 | 1.457 |  |  |  | -0.165 | 0.144 |
| conditionTrue:storyWeapon | 0.419 | 0.695 | -1.055 | 1.880 |  |  |  | 0.143 | 0.687 |
| speciesChimp:storyTool | -0.411 | 0.600 | -1.696 | 0.818 |  |  |  | -0.511 | -0.239 |
| speciesChimp:storyToy | -0.149 | 0.609 | -1.367 | 1.045 |  |  |  | -0.345 | 0.041 |
| speciesChimp:storyWeapon | -0.060 | 0.599 | -1.296 | 1.191 |  |  |  | -0.169 | 0.092 |
| conditionTrue:speciesChimp:storyTool | 0.761 | 0.847 | -0.986 | 2.478 | 1.488 | 3 | 0.685 | 0.513 | 0.992 |
| conditionTrue:speciesChimp:storyToy | 0.139 | 0.857 | -1.786 | 1.888 |  |  |  | -0.062 | 0.253 |
| conditionTrue:speciesChimp:storyWeapon | -0.215 | 0.835 | -1.926 | 1.524 |  |  |  | -0.493 | 0.075 |

Indicated are fixed effect estimates together with their standard errors, 95% confidence limits, significance tests and the range of estimates (min and max) obtained when excluding levels of random effect factors one at a time. All factors were dummy coded; reference categories were False for condition, bonobo for species, snake for story, and Edinburgh for location. Order was z-transformed to a mean of zero and a standard deviation (sd) of one, mean and sd of the original variable were 4.19 and 2.27.

**Table S8** Sample sizes and dispersion parameters.

| area | response | N total | N participants | dispersion parameter |
| --- | --- | --- | --- | --- |
| entire | pupil dilation | 140 | 30 | - |
| entire | prop. looking | 206 | 31 | 1.01 |
| box | prop. looking | 206 | 31 | -1.43 |

Note that ‘entire’ refers to the area of interest containing the protagonist and the box.
